# Mesenchymal stem cell suppresses the efficacy of CAR-T toward killing lymphoma cells by modulating the microenvironment through stanniocalcin-1

**DOI:** 10.1101/2022.09.21.508926

**Authors:** Rui Zhang, Qingxi Liu, Sa Zhou, Hongpeng He, Mingfeng Zhao, Wenjian Ma

## Abstract

Stem cells play critical roles both in the development of cancer and therapy resistance. Although mesenchymal stem cells (MSCs) can actively migrate to tumor sites, their impact on CAR-T immunotherapy has been little addressed. Using an in vitro cell co-culture model including lymphoma cells and macrophages, here we report that the CAR-T cell mediated cytotoxicity was significantly inhibited in the presence of MSCs. MSC caused an increase of CD4+ T cells and Treg cells but decrease of CD8+ T cells. In addition, MSCs stimulated the expression of indoleamine 2,3-dioxygenase (IDO) and programmed cell death-ligand 1 (PD-L1) that contribute to the immune-suppressive function of tumor. Moreover, MSCs suppressed key components of NLRP3 inflammasome by modulating mitochondrial ROS release. Interestingly, all these suppressive events hindering CAR-T efficacy could be abrogated if the STC1 gene, which encodes the glycoprotein hormone staniocalcin-1, was knockdown in MSC. Using xenograft mice, we confirmed that CAR-T function could also be inhibited by MSC in vivo and STC1 played a critical role. These data revealed a novel function of MSC and staniocalcin-1 in suppressing CAR-T efficacy, which should be considered in cancer therapy and may also have potential applications in controlling the toxicity arising from excessive immune response.

## Introduction

Advances in chimeric antigen receptor modified T cell therapy (CAR-T) in recent years has shown enormous promise in cancer immunotherapy, which has produced unprecedented clinical outcomes, most notably for patients with hematologic malignancies [1,2]. In spite of the striking achievements, CAR-T therapy are also facing many challenges such as the treatment-related severe toxicity and side effects, including cytokine release syndrome (CRS) and neurotoxicity [3,4]. CRS is the most common acute toxicity associated with excessive immune response that causes fever, hypotension, and respiratory insufficiency. The neurotoxicity induced by CAR-T therapy exhibits a diverse array of neurologic symptoms such as tremor, expressive aphasia and impaired attention. The precise mechanism that causes these life-threatening side effects remains unclear [4,5]. On the other hand, the success of CAR-T therapy in treating solid tumors is still very limited [6]. Identifying hurdles and potential mechanisms that impede the function of CAR-T cells is of vital importance in order to expanding its use. The immunosuppressive tumor microenvironment (TME) is one of the obstacles that diminish the efficacy of CAR-T therapy especially for solid tumors.

Among the many factors that can modulate TME and immune response, the impact of mesenchymal stem cell (MSC) on CAR-T therapy has been little studied. MSC is a type of adult stem cells with high proliferative activity and multidirectional differentiation capacity. However, MSCs have additional paracrine effects that are believed to be underlie their therapeutic functions[7]. By secreting a variety of cytokines into tissue microenvironment, it has been known that MSCs can modulate extracellular matrix, promote angiogenesis, suppress inflammation and apoptosis [8–10]. Some MSC-secreted cytokines, such as stromal cell-derived factor 1 (SDF-1) and stem cell factor (SCF), play important roles in hematopoietic and immune regulation [11,12]. In addition, studies suggest that MSCs can modulate the function of monocytic lineages cells, especially macrophages [13–15]. Some reports also showed that MSCs could directly affect the functionality and cellular responses of T cells, Tregs and memory T cells [16–18].

It was reported that hMSCs could be activated by LPS-stimulated macrophages to increase the expression and secretion of stanniocalcin-1 (STC1) [19]. STC1 was a mitochondria related glycoprotein originally identified as a calcium/phosphate regulating hormone in bony fishes, and later on it was found to be a pleiotropic factor involved in various degenerative diseases such as ocular and renal disease, as well as idiopathic pulmonary fibrosis [20,21]. STC1 could improve the cell survival and regeneration of MSCs in a paracrine fashion [22]. There were also evidences suggesting that STC1 played an oncogenic role in various type of tumors [23,24]. Based on a retrospective study of ~1500 clinical samples, it was concluded that high STC1 expression is associated with the poor clinical outcome of breast cancer [25].

Considering the pleiotropic role of STC1, especially its intercellular linkage between MSCs, cancer cells and macrophage simulation, it is interesting to know what role it plays in connection to the functions of MSC in TME. Therefore, we generated stable STC1 knockdown MSC cell line. With a cell co-culture model containing CAR-T cells, hMSCs, macrophages and Pfeiffer lymphoma cells to partially mimic tumor microenvironment together with a xenograft mice model, here we studied the impacts of MSC on CAR-T efficacy and the potential immune response change in the presence and absence of STC1.

## Materials and methods

### Cell culture and isolation of primary cells

HEK-293T (ATCC) was grown in DMEM (Gibco) supplemented with 10% FCS (Gibco). Peiffier cells were grown in RPMI 1640 medium supplemented with 10% FCS. human umbilical cord blood-derived MSCs (UCB-MSCs) were established from consenting mothers and processed within the optimal period of 6 hours as described [26]. Peripheral blood samples were obtained from healthy male donors (n = 3) in Tianjin First Central Hospital (clinical trial# ChiCTR-ONN-16009862). The scFv targeting CD19 plasmid was originated from the FMC63 clone. The CAR vectors containing scFv, human 4-1BB and CD3z signaling domains were subcloned into the pCDHMND-MCS-T2A-Puro lentiviral plasmid. The CAR sequence was preceded by the RQR8 tag separated by a short T2A peptide for detection purpose [27].

### Lentivirus Production

Preparation of the lentivirus was performed according to the manufacturer’s instructions (GeneCopoeia). Briefly, HEK-293T lentiviral packaging cells in DMEM supplemented with 10% heat inactivated fetal bovine serum followed by transfection when cells are 70-80% confluent. Dilute 2.5μg of lentiviral expression plasmid and 2.5 μg of Lenti-Pac HIV mix into 200μl of Opti-MEM® I (Invitrogen). In a separate tube diluting 15 μl of EndoFectin Lenti into 200μl of Opti-MEM I, then drop-wise adding to the plasmid mix and incubate for 10-25 minutes at room temperature. Collect the pseudovirus-containing culture medium 48 hours post transfection followed by ultracentrifugation and the pellets were resuspended in complete X-Vivo15 media and stored at −80°C until use.

### Production and detection of CAR-T Cells

CD3^+^ T cells from healthy donors were separated from PBMCs using CD3 immunomagnetic beads (#130-097-043, Miltenyi Biotec, Germany), then amplifing using CD3/CD28 stimulation beads (#11131D, Thermo Fisher Scientific) and IL-2 (100 IU/mL; Miltenyi Biotec) in X-VIVO 15 medium (Lonza). Cells were activated and expanded for 48 hours followed by transduction 2 hours later with lentivirus. T cells were generally engineered for 9-12 days to express a CD19-specific CAR, and stained with Alexa-Fluor 647-labeled polyclonal goat anti-mouse IgG (H+L) antibodies (Affinity) to detect CAR-T cells. All cells were further confirmed by staining with fluorescein isothiocyanate (FITC)-labeled anti-CD3 antibodies (Abcam).

### Cell co-culture model

THP-1 cells (5×10^5^/well) were seeded into six-well plates and polarized into M2 macrophages. After washing to remove all PMA and cytokines, the macrophages were cocultured with CAR-T cells and Preiffier cells. The ratio of CAR-T cells, tumor cells, and macrophages was 1:3:1.

### Generation of STC1 knockdown cells

Lentiviral particles PLKO.1 and PLKO.1-shSTC1 were provided by Beijing Institute of Radiation Medicine. Viruses were packaged by co-transfection with PLKO.1 and PLKO.1-shSTC1 into 293T cells. The supernatants containing viruses were collected 48h after transfection, then the centrifuged and resuspended lentivirus were used for further transduction of hMSCs in Opti-MEM. The stable STC1 knockdown hMSCs were obtained after 7-10 days of puromycin selection in 96-well plates. Transduction efficiency was determined by fluorescent microscopy.

### MTT assay

Cell viability was examined by 3-(4,5-dimethylthiazol-2-yl)-2,5-diphenyltetrazolium (MTT) assay (Sigma). The absorbance was measured using a Synergy™ 4 plate reader (Bioteck) with a test wavelength at 490 nm and a reference wavelength at 630 nm.

### Cell Migration determination by wound healing and transwell chamber assay

hMSCs^shCtl^ and hMSCs^shSTC1^ were grown in 6-well plates, and wounded using a sterile pipette tip. The progress of migration was recorded immediately following injury and photo-micrographs were taken at zero and 48 h.

For transwell assay, hMSCs^shCtl^ and hMSCs^shSTC1^ were seeded into the upper chamber of a transwell cell culture insert with 1.0×10^4^ cells in 200 μL of a 1% FBS-containing medium. The lower chamber was filled with 600 μL of medium containing 10% FBS. Twenty-four hours later, cells that had migrated to the lower side of the membrane were fixed in 4% paraformaldehyde and stained with DAPI. The migrated cells was counted and photographed in five fields of view, and was done in three independent experiments.

### Apoptosis detection with annexin V-FITC and PI and TUNEL assay

An increase in the plasma membrane phosphatidylserine (PS) externalization occurs early in apoptosis and can be detected by annexin V staining. hMSCs^shCtl^ and hMSCs^shSTC1^ were isolated and stained with annexin V-FITC and PI (Invitrogen), then apoptosis-positive cells were analyzed using FACS (Millipore Muse).

The terminal deoxynucleotidyl transferase-mediated dUTP nick-end labeling (TUNEL) assay was used to monitor the extent of DNA fragmentation as a measure of apoptosis [28]. After hMSCs^shCtl^ and hMSCs^shSTC1^ were fixed by formaldehyde, immunohistochemical detection of apoptotic cells was carried out using DeadEnd™ Fluorometric TUNEL System (Promega). The cells were washed with PBS and blocked with 10% goat serum, then using DAPI to stain nuclei. The samples were photographed with confocal laser microscope (Olympus), and TUNEL-positive cells was quantitated.

### Quantitative real time PCR (qRT-PCR)

Total RNA was extracted using TRIzol reagent (Invitrogen), serving as template for Real-Time PCR using random primers and M-MLV reverse transcriptase. The primers used were as follows: human TSP1: forward: 5’-TTGTTAAGAGGTTTGAGTAGGAGAG-3’ and reverse: 5’-CCCACCTTACTTACCTAAAATCACA-3’.

### Western blotting and cytokine release analysis

Western immunoblotting was performed as previously described [29]. After SDS-PAGE and blotting proteins were detected using the following antibodies: rabbit anti-IL-1β (Abcam, ab9722), anti-Caspase-1 p20 (Bioss, bs-10442R), AIM2 (Abcam, ab93015), IDO (Bioss, bs-15493R), PD-L1 (Bioss, bsm-54472R) and mouse anti-GAPDH (Santa Cruz) primary antibodies. The secondary antibodies were IRDye-800-conjugated anti-mouse and anti-rabbit immunoglobulin G (Li-COR Biosciences) (1:200). Immunofluorescence was detected using Odyssey Infrared Imaging System (Gene Company Ltd.). GAPDH expression was used as an internal control. The relative quantification of protein expression was analyzed using ImageJ software. The level of IL-1β in the serum was detected using ELISA by electrochemiluminescence (R&D Systems, France).

### Flow Cytometry

The expression of CD4、CD8、CD127 and CD25 in CAR-T cells was analyzed using flow cytometry with the following fluorochrome-conjugated monoclonal/polyclonal antibodies (all from Caprico Biotechnologies): anti-human CD4 (CD004210403), anti-human CD8 (CD008210301), anti–human CD127 (CD127210501), anti–human CD25 (CD025210301).

### in vitro analysis of CAR-T Cytotoxicity toward pfeiffer cells

Seeding CD19 CAR-T cells (4×10^5^ cells/group) in a co-culture with pfeiffer cells and macrophages polarized from M-THP1 at a 4:1 ratio and incubate for 48 h. The cell killing of CAR-T toward pfeiffer cells was determined using a lactate dehydrogenase (LDH) cytotoxicity test kit (Dojindo Molecular Technologies, Inc.) and measured at 0, 24 and 48 h after cell co-culture.

### Cellular and mitochondrial ROS detection

ROS was measured using CellROX Deep Red Reagent (Invitrogen) and MitoTracker Green FM Dye (Invitrogen) [30,31]. Briefly, cells co-culture for 24 hours followed by loading with CellROX dye (5 mM) and MitoTracker Green dye (100 nM) at 37 C for 30 minutes, then analyzed by flow cytometry. The data were analyzed using Flowjo software (Tree Star Inc., Ashland, OR)

### Xenograft tumor model

Female 6-8-week-old NOD/Shi-scid IL-2Rγ(null) (NOG) mice weighing 20±1.6 g (n=36, Vitonlihua Experimental Animal Technology Co., Ltd, Beijing, China) were injected with 5×10^6^ Pfeiffer cells expressing luciferase by subcutaneous injection on each side. All animal experiments were approved by the Ethics Committee of Tianjin First Central Hospital (ChiCTR-ONN-16009862). Established tumors were monitored by bioluminescence imaging (BLI). Upon confirmation of engraftment after 25 days, the mice were randomized into 3 groups and treated by tail vein injection of 5×10^6^ CD19 CAR-T cells and 2.5×10^6^ M-THP1. At the same time, 5×10^6^ cells/mice of hMSCs^shSTC1^ or hMSCs^shCtl^ were injected to multi-points of the tumor area. Tumor progression were photographed with BLI following intraperitoneal injection with D-luciferin (Goldbio, 150 mg/kg) at 14, 28 and 38 days. All the mice were sacrificed when either experimental or humane endpoints were reached.

### Immunohistochemical analysis of IL-1β in vivo

Mice were sacrificed at day 10 after CAR-T/M-THP1 and hMSC injection and tumor samples were fixed with formalin and embedded in paraffin. Tumor tissues were examined by immunohistochemistry staining as previously described [32]. Briefly, the sections were exposed to 3% H_2_O_2_ in methanol after deparaffinization and rehydration and then blocked with 1% BSA for 30 min at room temperature. After blocking, the sections were incubated with primary antibody (anti-IL-1β) overnight at 4°C, followed by incubation with peroxidase-conjugated secondary antibodies. IL-1β+ cells were quantified by measuring the number of stained cells.

### Statistical analysis

SPSS 17.0 (SPSS, Inc., Chicago, IL, USA) software was used for statistical analysis. Data are expressed as the mean ± standard error, analyzed by t-test with q-test for pairwise comparison. P<0.05 was considered as statistically significant difference.

## Results

### Stable knockdown of STC1 in hMSC inhibited cell migration, slightly suppressed cell proliferation, but no increase on apoptosis

To study the function of stanniocalcin-1, we first generated a stable knockdown cell line by lentivirus-based shRNA for STC1 gene, and the expression of STC1 protein was evaluated by Western Blot (Fig. 1A). STC1 stable knockdown in hMSCs exhibited a minor effect in cell survival (Fig. 1B) and slightly reduced proliferation rate based on the small increase in proportion of cells in G0/G1 phases versus that in S phase (Fig. 1C) as determined by MTT and FACS analysis. To investigate whether knockdown of STC1 affect cell migration, wound healing and transwell chamber assays were performed. After creating a “scratch” in a monolayer of hMSCs, the closure of the gap was determined after 24h. As shown in Fig. 1D, comparing to control hMSCs, the gap was evidently less filled in hMSC^shSTC1^. The inhibitory effect on cell migration was further confirmed by transwell assay. As shown in Fig 1E, there was significant migration and invasion observed in hMSCs^shCtl^, whereas there was a >30% reduction in migration across the transwell chamber membrane in hMSCs^shSTC1^. To further determined whether knockdown of STC1 may have any lethal effect, apoptosis was determined by two different assays. To measure early apoptosis, cells were stained with the Alexa Fluor® 488 annexin V and PI followed by flow cytometry to detect apoptosis-associated phosphatidyl serine expression and membrane permeability (Fig. 1F). Parallelly, DNA fragmentation was determined with TUNEL assay (Fig. 1G). Both studies showed that knockdown of STC1 did not cause apoptosis of hMSCs.

**Fig. 1.**
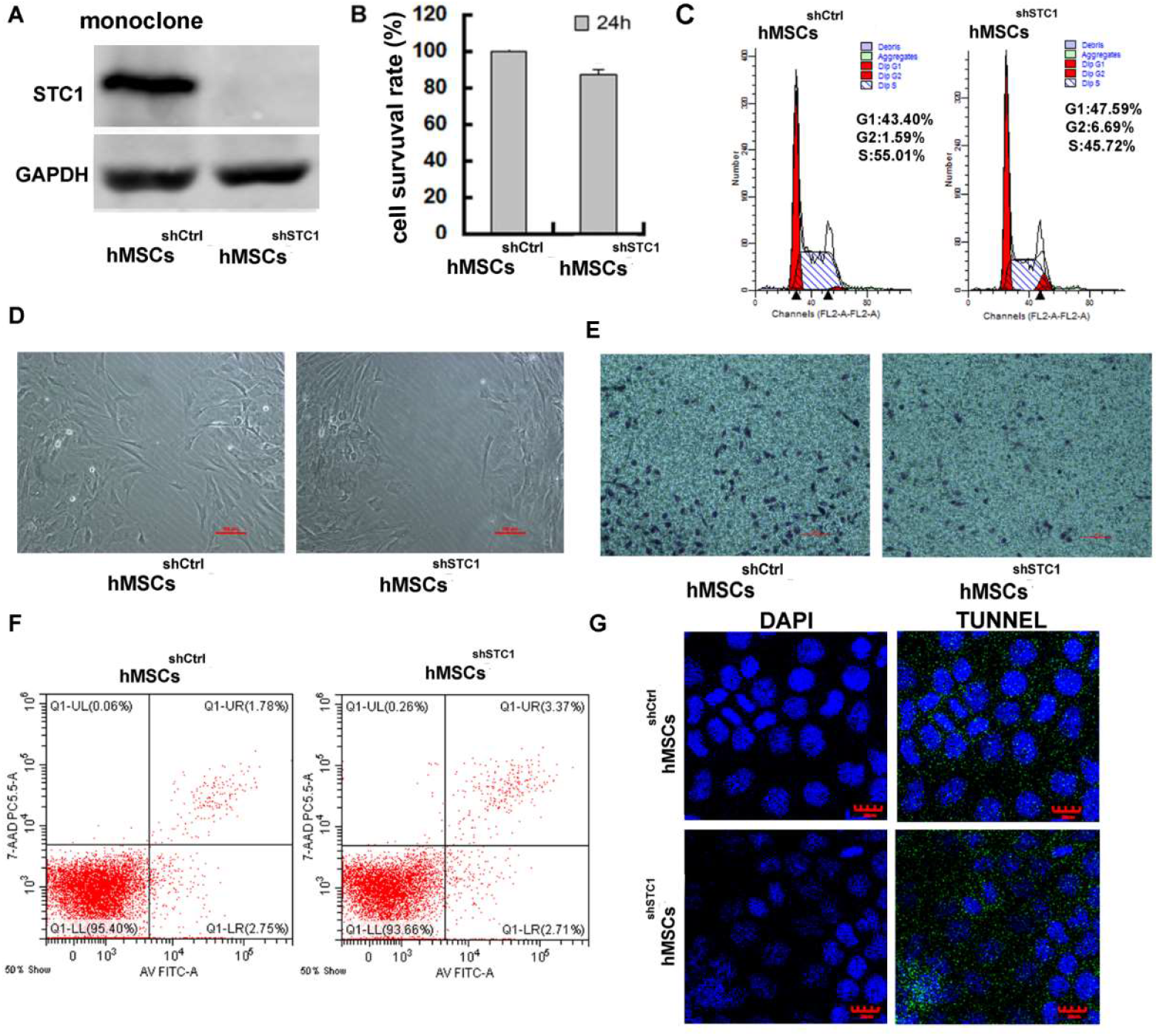
The impact of STC1 knockdown on cell proliferation, migration and apoptosis of hMSCs. A) Western blot of STC1 protein expression in hMSCs. B) cell viability determined by MTT. C) FACS analysis of cell cycle progression. D, E) knockdown of STC1 suppressed cell migratory as determined by wound healing and transwell chamber assays. F) apoptosis analysis by the Alexa Fluor® 488 annexin V and PI detection. G) DNA fragmentation determination by TUNEL.

### The presence of hMSCs inhibited CAR-T cell killing activity, but knockdown of STC1 completely abrogated this inhibition

To investigate the impact of hMSCs on CAR-T treatment, we used an in vitro cell co-culture model modified according to previous studies to mimic a simplified situation of tumor environment [33,34]. The co-culture contained CD19 CAR-T cells, Pfeiffer cells that were from human diffuse large cell lymphoma, and M2 macrophages (derived from THP-1 cells by PMA polarization for 24 hours) at cell number ratio 1:3:1. The cell killing activity of CAR-T cell toward Pfeiffer cells were determined by LDH cytotoxicity assay. As shown in Fig. 2A, 67% of pfeiffer cells were killed after being exposed to CAR-T cells for 24 hours, and 93% were killed at 48 hours. After adding hMSCs into the co-culture, the cell killing activity of CAR-T was significantly inhibited (fig. 2A). The number of hMSC added was the same as CAR-T cell. Interestingly, the inhibitory effect of hMSCs on CAR-T cytotoxicity could be completely abrogated if knockdown STC1 gene in hMSCs. These results for the first time revealed that CAR-T efficacy could be affected by the presence of MSCs and the gene STC1 played a critical role.

**Fig. 2.**
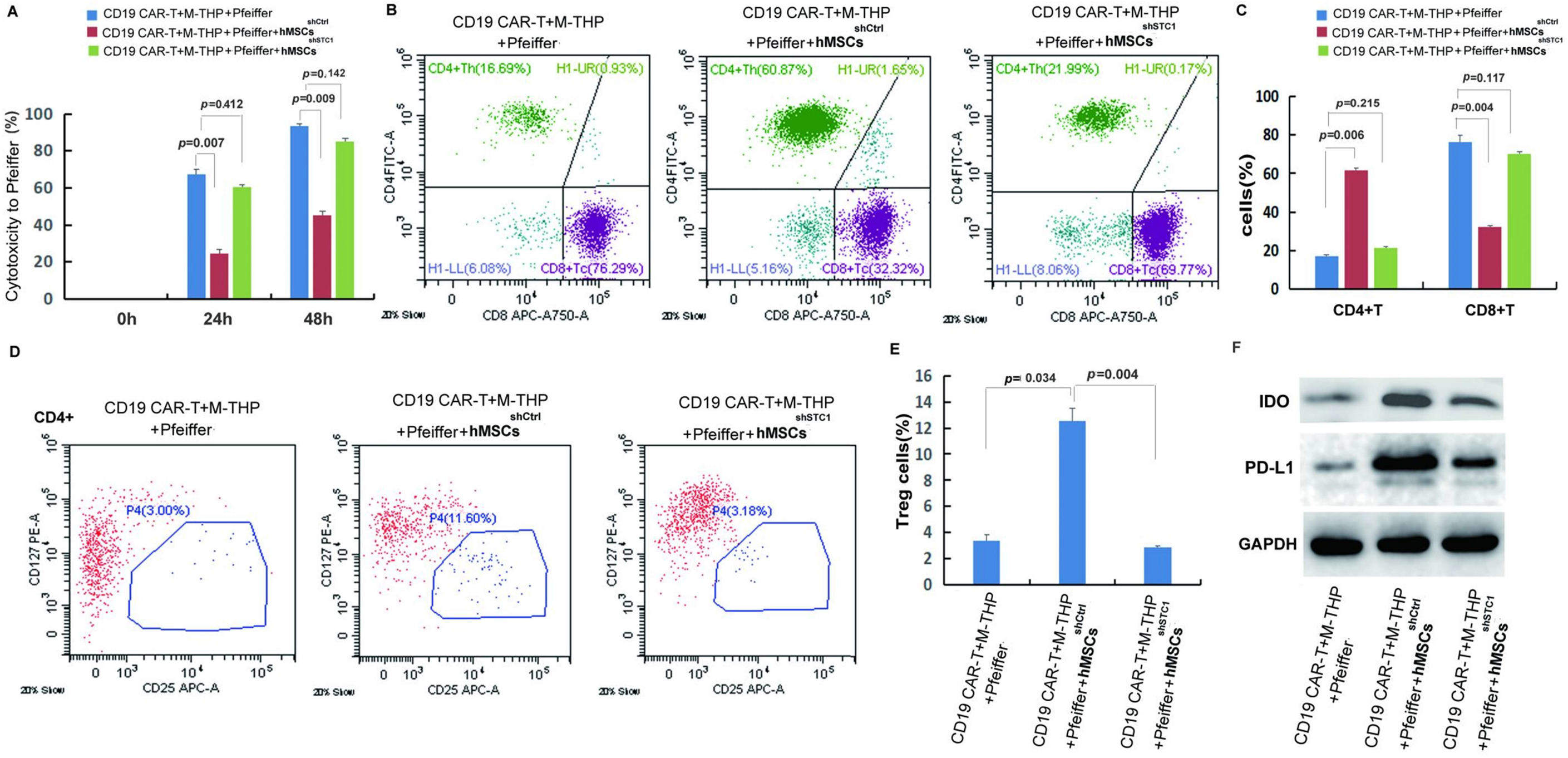
Analysis of cytotoxicity, T cell composition and immune suppressive markers. A) hMSC induced CAR-T cytotoxicity change in the presence and knockdown of STC1 gene; B) FACS analysis of CD4^+^ and CD8^+^. C) Quantitation of CD4^+^ and CD8^+^; C) FACS analysis of Treg^+^ cells (CD4^+^CD127^+^CD25^+^); D) Quantitation of Treg+ cells. E) Western blot analysis of IDO and PD-L1. (Please see next page for Fig.2)

### Co-culturing with hMSCs caused an increase of CD4+ T cells and Treg cells but decrease of CD8+ T cells

Previous studies have demonstrated that the composition of CD4+ and CD8+ T cell subsets was crucial for CAR-T cell efficacy [35,36]. To investigate the mechanism on how hMSC inhibited the cytotoxicity of CAR-T, the amount of CD4+ and CD8+ T cells were analyzed by flow cytometry at 24 hours after co-culture. As shown in Fig. 2B and 2C, the ratio between CD4+ and CD8+ was about 1:4 when there were no hMSCs in co-culture (Fig. 2C). However, the addition of hMSC caused significant increase of CD4+ and decrease of CD8+ T cells (Fig. 2B), resulting in a ratio change to 2:1. Similar to the change of CD4+ T cells, the percentage of regulatory T cells (Treg) was also significantly increased from ~3% to 12% when co-culture with hMSC (Fig. 2D and 2E). When using hMSC^shSTC1^, all the changes were completely reversed back to the level similar to that of co-culture without hMSCs. This explains the reduced CAR-T cytotoxicity since CD8+ T cells are directly responsible for specific lytic activity against lymphoma [35]. Tregs, which account for 5-10% of the total number of CD4+ T cells, are known to play a role in suppressing the function of T cells and other immune cells [37]. Therefore, the above results indicate that hMSCs’ inhibitory effect on CAR-T cytotoxicity was due to both suppression of CD8+ cells and the induction of Treg cells, and the presence of STC1 was indispensable for these impacts of hMSC.

### The presence of hMSC enhanced immune suppression and STC1 played a key role

The immune-suppressive TME is the main cause of CAR-T cell exhaustion which attenuates its efficacy. To further investigate the function of STC1 and the molecular mechanism of hMSC on CAR-T resistance, some key regulators of TME were determined. As shown in Fig. 2F, the addition of hMSC to the cell co-culture stimulated the expression of indoleamine 2,3-dioxygenase (IDO) and programmed cell death-ligand 1 (PD-L1). IDO and PD-L1 are two of the most important immunosuppressive proteins. IDO is an intracellular enzyme that converts tryptophan into inhibitory metabolites for T - cell activity [38]. PD-L1 is expressed in tumor cells and immune cells contributing to the immune-suppressive TME [39]. When using hMSC^shSTC1^, the expression level of IDO and PD-L1 were both significantly reduced by more than 50%, though still higher than that of without hMSC. These results suggest that presence of hMSC can enhance the expression of immune suppressive proteins in pfeiffer cells and macrophages, and the presence of STC1 is important for hMSC to exert these effects.

### hMSCs suppressed key components of NLRP3 inflammasome by modulating mitochondrial ROS release

In the co-culture model, M2 macrophages were included since previous study showed that macrophages could activate hMSCs to secrete STC1 [16]. In addition, macrophage is a critical part of immune response and important regulator of immunotherapy [40]. To further identify the mechanisms mediating the inhibitory effects of hMSCs, the activation of NLRP3 inflammasome were determined. The NLRP3 inflammasome is a critical component of the innate immune system mediating caspase-1 activation and proinflammatory cytokines secretion in response to harmful stimuli such as infection and endogenous stress [41]. As shown in Fig 3A, the release of cleaved caspase-1 p20 in cell lysates, which is the indicator of caspase-1 activation, was detected after PMA polarization of M-THP. Following co-culture with CD19 CAR-T, the level of cleaved caspase-1 was significantly upregulated. The increase of active caspase-1 was evidently abrogated when hMSCs were added into the co-culture. knock-down of STC1 led to another reverse and completely blocked the inhibitory function of hMSCs (Fig. 3A). Concomitant with the reduction in active caspase-1, the cleaved IL-1β mature form and absent-in-melanoma 2 (AIM2), two key components of the inflammasome [42], were both increasingly expressed following M-THP polarization and further incubation with CAR-T (Fig. 3A). Comparing to the partial inhibition of the active caspas-1 formation, addition of hMSC in the cell co-culture showed a stronger inhibition of these two proteins and their expression level were returned back to base level of Pfeiffer plus CAR-T (Fig. 3A). This result suggests that the immune suppressive effect of hMSC was through its impact on macrophages, not CAR-T or Pfeiffer cells. Knockdown of STC1 abrogated the inhibition of hMSC on IL-1β and AIM2 (Fig. 3A). The levels of IL-1β in the supernatants measured by ELISA showed similar results as cell lysate (Fig. 3B).

**Fig. 3.**
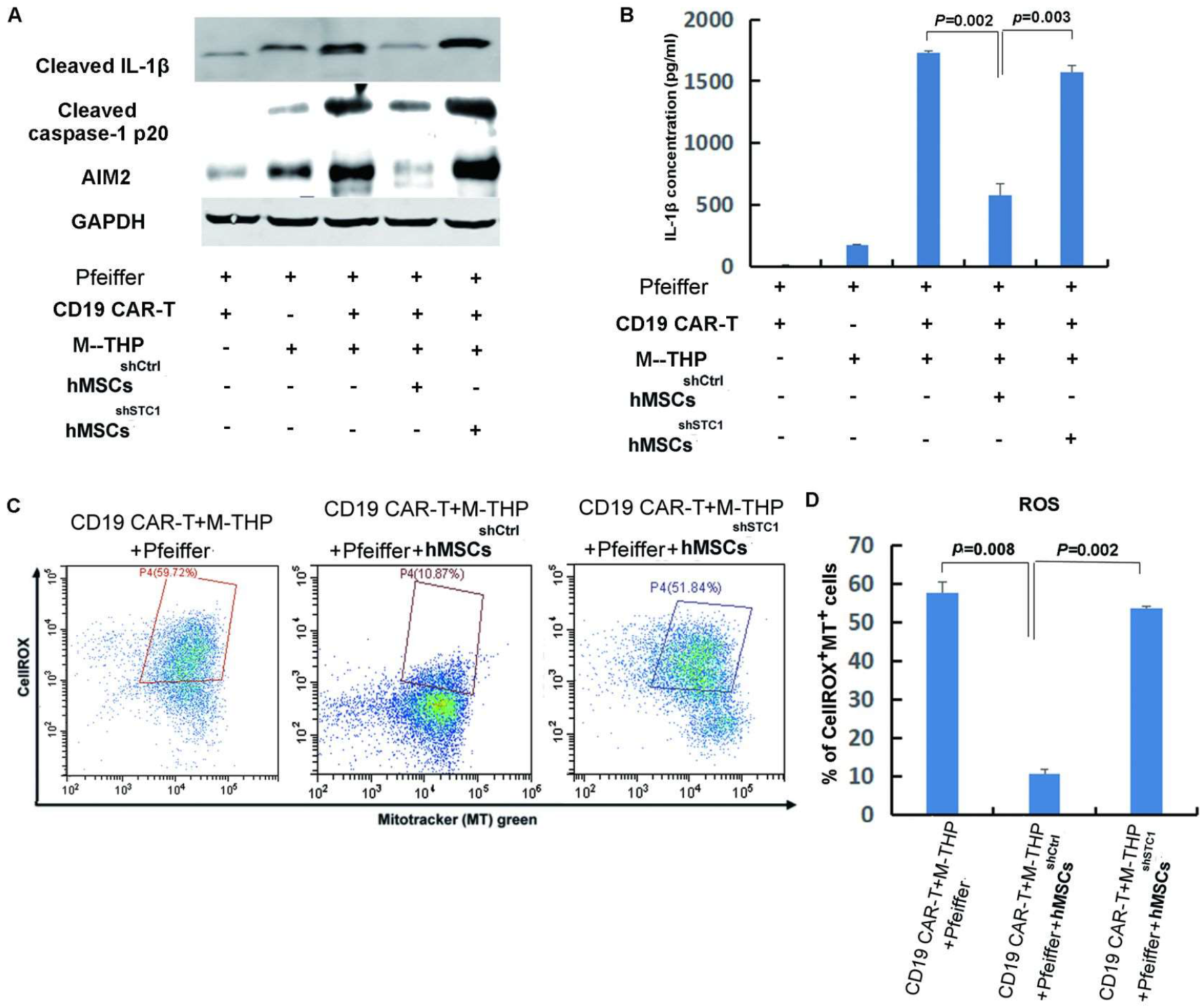
The impact of MSCs on the expression of key components involved in the formation of NLRP3 inflammasome and mitochondrial ROS. A) The protein expression of IL-1β, caspase-1, and AIM2 in cell lysates analyzed by Western blot. B)Quantitation of IL-1β secretion in the supernatants by ELISA. C) FACS analysis of ROS level and mitochondria mass with fluorescent dye CellROX Deep Red and MitoTracker Green. D) Quantitation of mitochondria specific ROS level based on the percentage of cells that were both positive for CellROX and MitoTracker.

Considering that mitochondrial dysfunction is one of the major stimuli that activate NLRP3 inflammasome, and it was reported that exogenous STC1 is internalized by macrophages within 10 min and localizes to mitochondria to suppresses superoxide generation [43]. We determined the impact of hMSC on the intracellular level of reactive oxygen species (ROS) and mitochondria mass in macrophage by fluorescent dye CellROX and MitoTracker Green, respectively. As shown in Fig. 3C and 3D, the presence of hMSCs^shCtl^ markedly suppressed both the cellular and mitochondrial ROS induced by the co-culture of CAR-T cells, tumor cells and macrophages. knockdown of STC1 eliminated the function of hMSC in suppressing ROS. This result correlates well with the expression of caspase-1, IL-1β and AIM, suggesting that hMSCs inhibited NLRP3 inflammasome activation in macrophages was most likely by inhibiting the oxidative burst.

### hMSCs showed strong inhibition on CD19 CAR-T therapy in xenograft mice, which was abrogated by STC1 knockdown

The immune suppressive impact of hMSC on CAR-T therapy and the function of STC1 was further evaluated in xenograft model. Upon injection of Pfeiffer cells and confirmation of engraftment, we injected hMSC into the tumor area while applying CAR-T treatment by tail vein injection. As shown in Fig. 4A, CD19 CAR-T treatment combined with injection of hMSC^shSTC1^ achieved a significant curative effect and the tumors nearly disappeared at day 38. However, hMSC^shCtl^ group showed continue increase in tumor size and spreading of tumor.

**Fig. 4.**
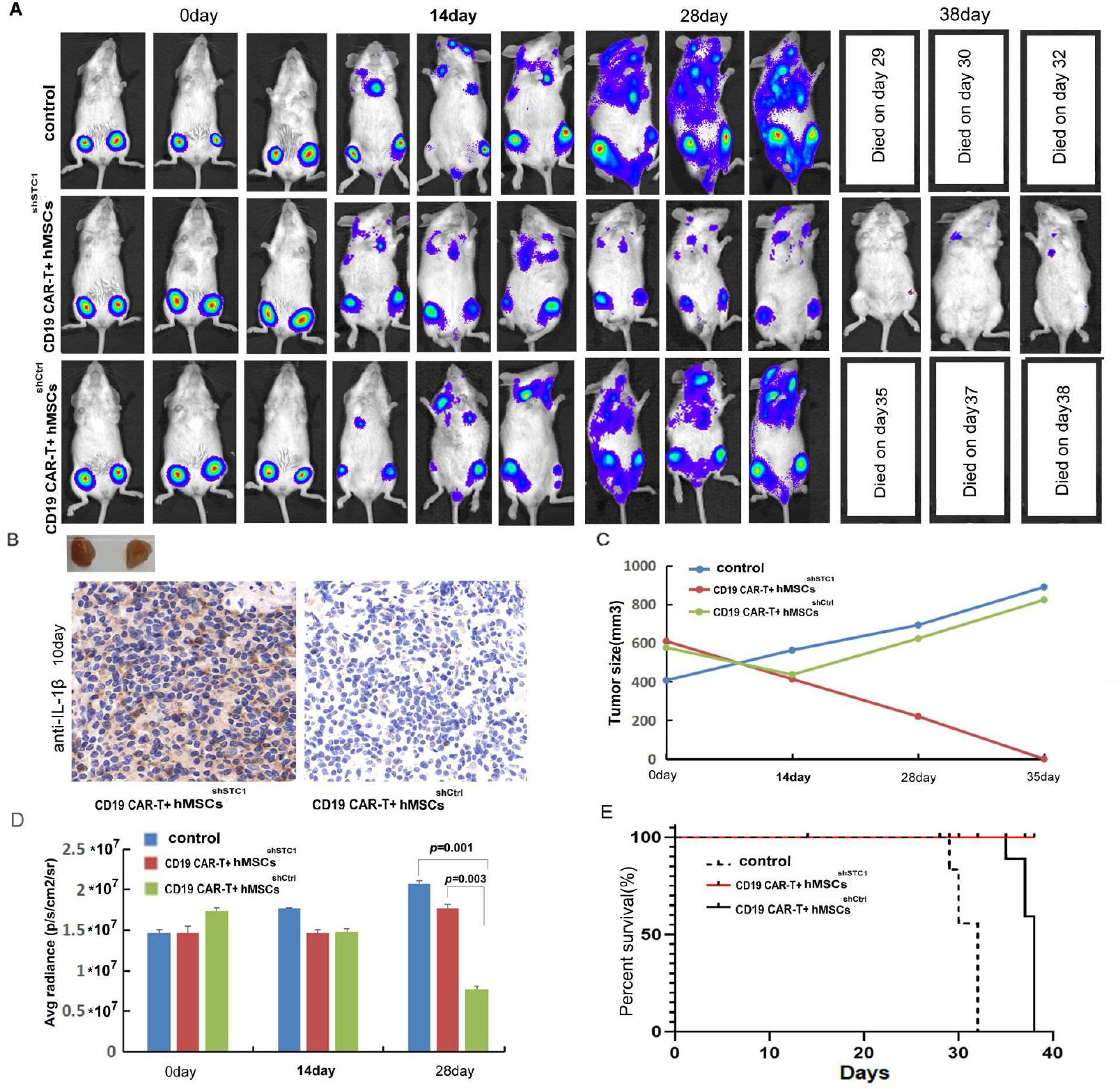
The inhibition of hMSC on CAR-T therapy in xenograft mice relyed on STC1. A) The formation and progression of tumor in three groups of mice monitored with bioluminescence imaging (BLI): The control group without any treatment, CAR-T/M-THP1/hMSCs^shSTC1^ group and CAR-T/M-THP1/hMSCs^shCtrl^ group. Day 0 was set when the engraftment was confirmed after injecting the Pfeiffer cells. B) immunohistochemical analysis of IL-1β in tumor tissue at day 10, positive cells display a brownish-yellow staining. C) The tumor size change with time. D) The counted average radiance; E) The survival rate of three mice groups.

Based on immunohistochemical analysis of IL-1β in tumor tissue at day 10, the number of positive cells (brownish-yellow staining) ranged from 76% to 100% in hMSC^shSTC1^ group, while it ranged from 5% to 20% in hMSC^shCtrl^ group, indicating that hMSC could suppress TME and STC1 knockdown significantly diminished this impact (Fig. 4B).

The changes of the average radiance were consistent with the changes of the tumor size (Fig. 4C and 4D). The survival time of mice demonstrated that mice in CAR-T combined with hMSC^shSTC1^ group had the longest survival with no death by day 38 (Fig. 4D). Comparing to the control group of no CAR-T treatment, tumor spreading in hMSC^shCtl^ group was slower and all survived for 6 days more. These results confirmed the inhibitory effects of hMSC on CAR-T therapy under in vivo situation, and demonstrated that STC1 is an important factor affecting therapy efficacy.

## Discussion

Stem cells are believed to play critical roles in resistance to cancer therapy, which is a major contributor to poor treatment responses and tumor relapse. Previous studies have been mainly focused on the role of cancer stem cells. In the current study we presented evidences that the presence of MSCs in TME may also be an important source of cancer treatment resistance. By modulating TME, MSCs showed a strong suppressive function on CAR-T efficacy toward lymphoma cells, and interestingly the presence of STC1 gene played a critical role.

The role of stanniocalcin-1 in cancer is paradoxical. Some reports showed that it exerts an oncogenic role, whereas other studies suggested the opposite [44]. The aberrant expression of STC1 have been reported to impact various types of cancer, such as triggering tumor angiogenesis by upregulating the expression of VEGF in gastric cancer cells [45], causing tumorigenesis and poor clinical outcomes in ovarian, colorectal and lung cancers [20,44]. To date the potential roles of STC1 in immunotherapy are still largely unknown. Here we demonstrated that the presence of STC1 is critical for MSC to exert its immunosuppressive role by inhibiting cytotoxic T cell subsets, activating some key immune suppressive/escape mechanisms and crosstalk with other immune cells.

First, a significant downregulation of CD8+ T Cells together with the upregulation of CD4+ T helper cell subsets and Tregs indicated that the suppressed CAR-T efficacy was at least partially associated with MSC’s function in modulating the proliferation of different T-cell subsets. Since the suppression of CD8+ T cells was completely abrogated if knockdown STC1 in MSCs, it is clear that STC1 played a key role here. Moreover, considering that STC1 is secreted into the extracellular matrix in a paracrine manner, MSCs’ modulation of the T cell subsets is most likely indirectly via altered cytokine expression or other secondary molecules activated by STC1.

The presence of MSCs also stimulated the expression of IDO and PD-L1, two important immune suppressive molecules. Upregulation of IDO is an endogenous feedback mechanism controlling excessive immune responses, which can be produced both by tumor cells and macrophages [47]. IDO-mediated formation of immunosuppressive metabolites can inhibit T-cell proliferation and induce T-cell death through the dioxin receptor [48,49]. PD-L1 is a well characterized molecule of the major escape mechanism to immunotherapy by inhibiting PD-1 mediated effector T cell function and downregulating antigen tolerance[39]. There have been numerous studies reporting the bidirectional interactions between MSCs and cancer cells, resulting in regulating the expression of PD-L1 in the surface of various cancer cells or TME [50–53]. Importantly, here we demonstrated that the upregulated expression of both IDO and PD-L1 by MSCs were much reduced if STC1 gene was knockdown.

The paracrine activity of MSCs is now widely recognized as an important cellular mechanism to communicate with immune cells and various other cell types in TME [54]. Consistent with previous studies, we found that addition of hMSCs to the co-culture cell model suppressed the formation of NLRP3 inflammasome in macrophage as determined by the downregulation of some key proteins including IL-1β, the activated caspase-1 and AIM. It was reported that CD4^+^ T cells could inhibit inflammasome-mediated caspase-1 activation and IL-1β release through TNF ligands or by interferon signaling [55]. Therefore, the modulation of T-cell subsets and activation of NLRP3 inflammasome by hMSC appears to be closely connected. Since NLRP3 inflammasome is a key factor in the neuroinflammation onset in CNS injuries [41], the suppression of NLRP3 inflammasome by hMSC may be potentially beneficial in reducing the exacerbated immune responses associated with CAR-T therapy.

The formation of NLRP3 inflammasome was reported to be through NF-κB dependent transcription of IL-1β, IL-18, and NLRP3, whereas its activation is triggered by extracellular stimuli such as lysosomal permeability, potassium efflux, and oxidative stress [42]. Considering that exogenous STC1 could be internalized by macrophages within 10 min and localizes to mitochondria and played suppressing role on ROS generation [43], we speculated that the inhibition of NLRP3 inflammasome formation might be a feedback mechanism occurred between macrophage and hMSC.

In fact, it was reported that LPS-stimulated macrophages do stimulate the expression and secretion of stanniocalcin-1 in hMSCs [19]. Our data further demonstrated that knockdown of STC1 deprived the function of hMSC in suppressing all the three markers used in the current study in determining NLRP3 inflammasome formation, as well as the suppression for mitochondria ROS production. These data support the idea that a feedback regulation mechanism exists between hMSC and macrophage during CAR-T therapy.

Using Xenograft mice model, we confirmed that the tumor killing efficacy of CAR-T could also be inhibited by hMSCs in vivo, whereas knockdown of STC1 effectively abolished the inhibition. Need to note that the amount of the injected hMSCs were much higher than that of the in vivo situation. Nevertheless, the results give clear indication that STC1 is critical for the immune suppressive function hMSC.

In summary, the present study revealed a significant impact of hMSC in suppressing CAR-T efficacy, and provided evidences that the STC1 gene played a critical role by the regulation of various immune suppressive mechanisms. A speculative schema of the signaling and interactions among hMSC, macrophage, CAR-T and tumor cell based on our current data is shown in Fig. 5. While further studies are needed to understand the detailed molecular interactions underlying, the findings we presented here is no doubt that would have potential clinical applications toward improving the efficiency of CAR-T therapy as well as reducing the excessive toxicity by modulating the level of STC1 in TME.

**Fig. 5.**
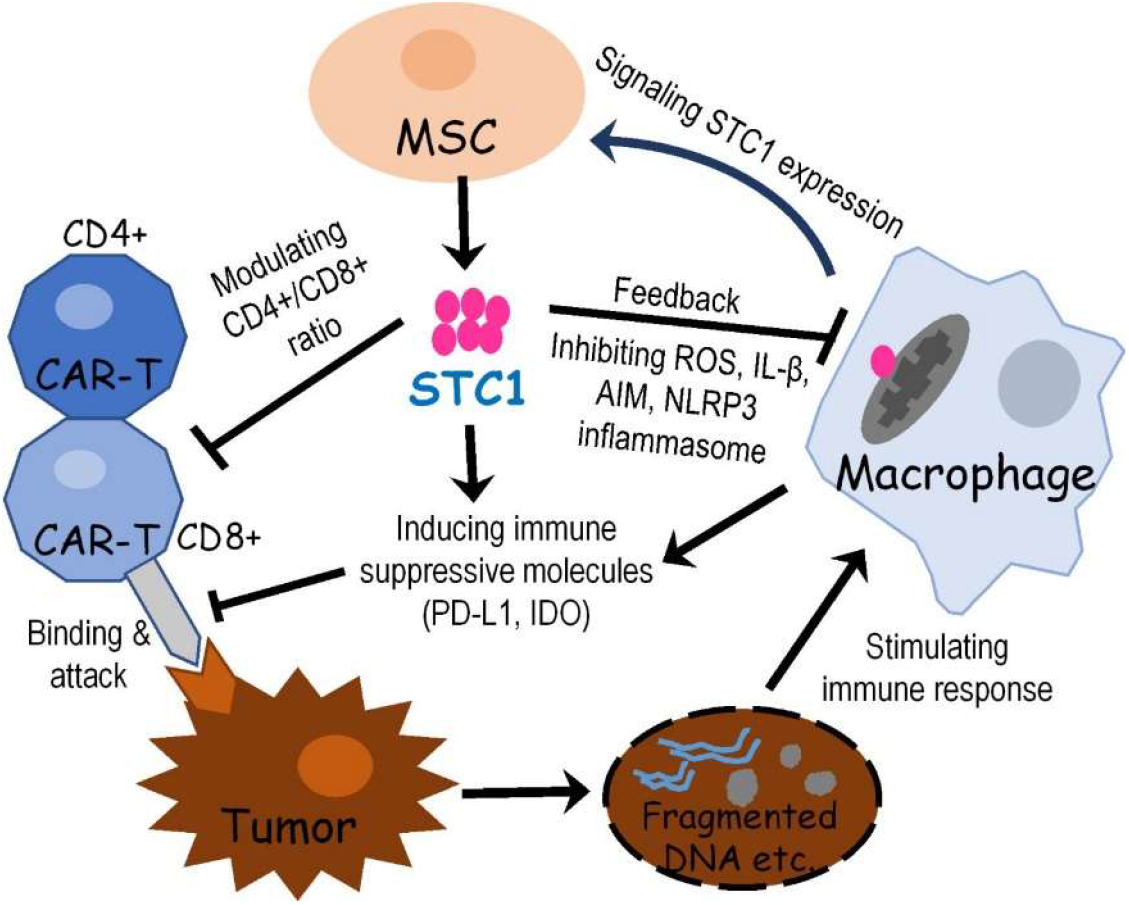
Proposed signaling and interactions among hMSC, macrophage, CAR-T and tumor cell. When cancer cells were destroyed by CAR-T cell, the release of fragmented DNA and other stimulating factors activated the release of mitochondria ROS and formation of NLPR3 inflammasome. Signals from activated macrophage and other extracellular molecules as well as oxidative stress may stimulate MSC to express and secrete STC1. Then STC1 serves as a pleotropic factor to suppress CAR-T cytotoxicity and other immune responses via direct or indirect pathways.

## Acknowledgments

This work was supported by the National Key R&D Program of China (2018YFA0901702), the Shandong Key R&D Program (2019GSF107088) and the National Science Foundation of Shandong (ZR2020MC077).

